# Intra-individual behavioural variability: a trait under genetic control

**DOI:** 10.1101/795864

**Authors:** Rie Henriksen, Andrey Hoeglund, Jesper Fogelholm, Robin Abbey-Lee, Martin Johnsson, Niels Dingemanse, Dominic Wright

## Abstract

When individuals are measured more than once in the same context they do not behave in exactly the same way each time. The degree of predictability differs between individuals, with some individuals showing low levels of variation around their behavioural mean while others show high levels of variation. This intra-individual variation in behaviour has received much less attention than between-individual variance in behaviour and very little is known about the underlying mechanisms that affect this potentially large but understudied component of behavioural variation. In this study, we combine standardized behavioural tests to estimate intra-individual behavioural variance with a large-scale genetical genomics analysis to identify genes affecting intra-individual variability in an avian population. Our study shows that within-individual variance in behaviour has a direct genetic basis which is largely unique compared to the genetic architecture for the standard behavioural measures they are based on. We identify six candidate genes underlying variation in intra-individual behavioural variability many of which have previously been linked to behaviour and mental health. These findings demonstrate that within-individual variability in behavioural is a heritable trait in and of itself on which evolution can act.

## INTRODUCTION

Individuals within a population often differ consistently in many aspects of their behaviour (Sih, Bell et al. 2004). Consistent differences in behavioural traits, such as aggressiveness, shyness, sociability and activity between individuals have given rise to the growing field of animal personality and has motivated the development of many evolutionary theories aimed at understanding the processes that allow and maintain such within population variation (Réale, Reader et al. 2007).

This consistency does not mean that the behaviour of an individual is readily predictable. An increasing number of studies have shown that when individuals are measured more than once in the same context they do not behave in exactly the same way each time (Bell, Hankison et al. 2009, Stamp et al., 2012). The degree of within-individual variation differs between individuals, with some individuals showing low levels of variation around their behavioural mean while others show high levels of variation (Cleasby, Nakagawa et al. 2015).

Whereas proximate and ultimate causes of inter-individual variation in behaviour has been an area of intense research interest across animal taxa (Roff, 2002, Van Oers et al., 2005, Coppens et al., 2010), within-individual variation in behaviour is often assumed to be homogenous across individuals. Therefore, not much is known about the factors that affect this large but understudied component of behavioural variation (Bell, Hankison et al., 2009). Insight into the potential genetic control of intra-individual behavioural variability is crucial for disentangling its proximate causes and thereby fine-tune tests of hypothesis about the evolution of this type of behavioural variance.

In this study, we combine standardized behavioural tests to estimate within-individual variation in anxiety and sociability related behaviours with large-scale genetical genomics analysis to identify genes affecting intra-individual behavioural variability in a population of chickens. By combining quantitative trait locus (QTL) and expression QTL (eQTL) analyses of the brains of an advanced intercross based on Red Junglefowl (the wild progenitor of the modern domestic chicken) and domestic White leghorn chickens we identify putative genes underlying phenotypic differences in intra-individual behavioural variability and to what degree these differ between behavioural trait.

## METHODS

### Chicken study population and cross design

The population used in this study was an eighth generation intercross between a population of Red Junglefowl, derived originally from Thailand and a line of selected White Leghorn chickens (Schütz *et al*. 2002, 2004), with a total of 572 F8 individuals used in this study. This advanced intercross has already been used to identify candidate genes underlying variation in anxiety and sociability related behaviours (Johnsson et al., 2016; Johnson et al. 2018). The birds were behaviourally tested between the age of 3 weeks and adulthood. The hypothalamus was dissected out at 212 days of age and RNA extracted. For further details on feed and housing see Johnsson *et al*. (2012). The study was approved by the local Ethical Committee of the Swedish National Board for Laboratory Animals. All methods were performed in accordance with the relevant guidelines and regulations.

### Behavioural phenotyping

#### Defining intra-individual variability as a trait

To test if intra-individual behavioural variability is trait-specific or can be viewed as a trait of its own that transcends different behaviours and situation we tested intra-individual variability in several behavioural traits across three separate anxiety related tests (Open field (OF), Social reinstatement (SR) and Tonic immobility (TI) – see below for more details). Within-individual variation in behaviour between trials was used to calculate the magnitudes of Intra-individual variation (IIV, see figure 1, and below). With increasing magnitude indicating increasing intra-individual variation. Initially the magnitude of intra-individual variation was calculated for each behaviour (IIV_trait_). Secondly, these calculations of magnitude were used to calculate an average intra-individual variation magnitude per test (IIV_average_) and finally a global magnitude of intra-individual variation was calculated for each individual based on all the recorded behaviours from the three tests (IIV_global_), see below for more details).

**Figure 1.**
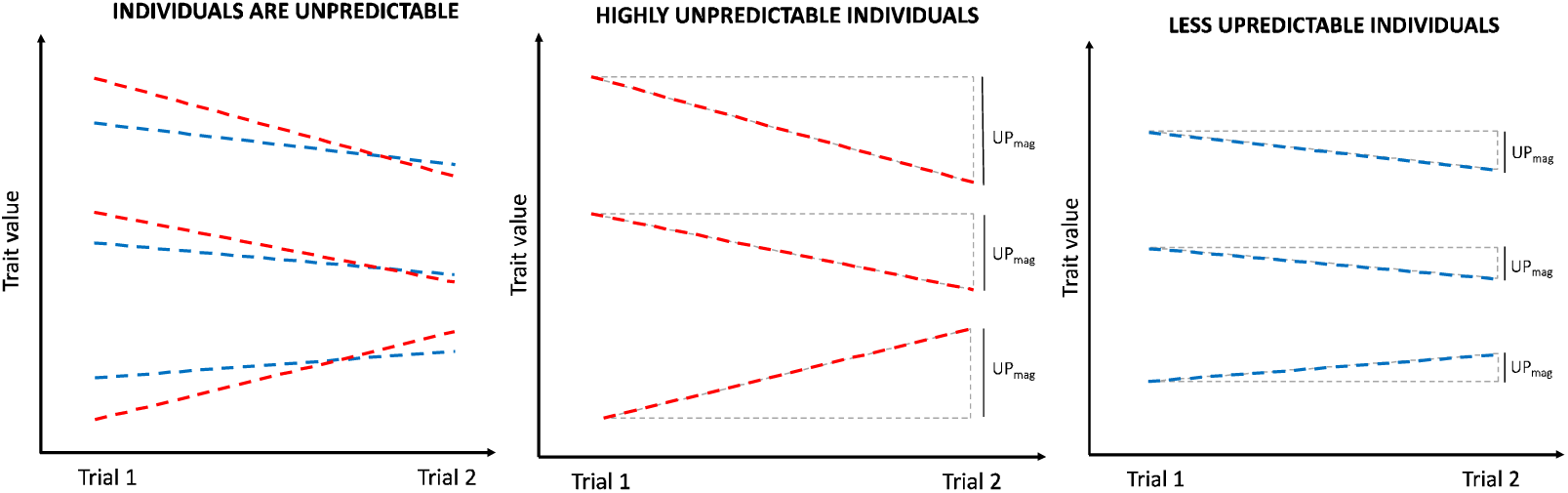
Illustration of intra-individual variation. The magnitude of intra-individual variation was calculated as the absolute difference between the behavioral trait value obtained during the first trial and second trial.

#### Calculating the magnitude of intra-individual behavioural variability

Intra-individual variation (IIV) was calculated for each behavioural trait (IIV_trait_) as the absolute difference in values obtained for that trait in trial 1 versus trial 2. Then an average intra-individual variation (IIV_average_) was calculated for each test situation as the sum of all IIV_trait_ calculated within that test situation. Finally, a global intra-individual variation (IIV_global_) was calculated for each individual based on all the recorded behaviours from the three tests.

IIV = Intra-individual variation

IIV_trait_ = |Trait _Trial-1_ – Trait _Trial-2_ |

IIV_average_ = (Sum of all IIV_trait_ within a test) / (Number of traits measured in test)

IIV_global_ = (Sum of all IIV_trait_ for all test) / (Total number of traits measured)

#### Social reinstatement

The social reinstatement test (Suarez and Gallup, 1983), measures social motivation and anxiety, with stressed chicks exhibiting a stronger social cohesion response (Marin et al., 2001). In this test, the individual is placed at one end of a narrow arena, with conspecifics located at the far end. The amount of time the individual spends associated with the conspecifics as opposed to exploring the remainder of the arena is considered a measure of sociality and anxiety. A more social or anxious animal will spend more time associating with conspecifics and will approach the conspecifics more rapidly, and therefore spend less time in the start zone of the arena (Marin et al., 2001). Trials were performed in a 100cm × 40cm arena. The social zone measured 20cm × 40cm and was adjacent to a wire mesh compartment containing three unfamiliar conspecific birds of the same age. Birds were placed in the start zone of the arena (also measured 20cm × 40cm) in the dark, prior to the lights being turned on and the trial beginning. Measurements were taken using the Ethovision software and continuous video recording (Noldus Information Technology, www.noldus.com). For each trial, total distance moved, velocity, length of time spent in the stimulus zone, latency to first enter the stimulus zone, and length of time in the start zone were measured. Each trial was five minutes and replicated twice per individual, with one week between an individual’s first and second test. Individuals were immediately removed from the arena upon the completion of the test to reduce potential habituation. Trials were replicated twice per individual, with each trial being 5 min in length. Trials were performed at 3 weeks of age. There was 1 week between an individual’s first and second trial. Individuals were immediately removed from the arena upon the completion of the test to reduce potential habituation

#### Open field

The open field assessment is a standard anxiety measurement, where the individual is placed alone in a brightly lit novel area after which behaviour is measured for a fixed duration and has been performed in a variety of vertebrates and invertebrates (Prut and Belzung 2003). In our study, trials were performed in a 100 × 80-cm arena at 4 weeks of age. Individuals were placed in the corner of the arena in complete darkness, prior to the test starting, with the lights turned on immediately at the commencement of the test. Trials lasted 5 min. Measurements were taken using the Ethovision software and continuous video recording (Noldus Information Technology, www.noldus.com). For each trial, total distance moved, proportion of time spent in the central zone (The 60 × 40-cm area in the middle of each arena was considered to be the central zone), velocity and frequency (number of times) that the central zone was entered were measured. Each trial was performed twice, with ∼1 week between an individual’s first and second trial. Individuals were removed from the arena immediately upon the test finishing to reduce potential habituation.

#### Tonic immobility

The third test, the tonic immobility test, measures an individual duration of immobility after being placed on its back and is thought to be a defence strategy evolved to reduce a predator’s interest in the prey, when the prey stops moving after it has been caught. The longer the animal stays in this immobile state the more fearful it is considered to be. In our study, the test bird was placed on its back in a V-shaped wooden cradle (approximately 50cm in length) and held by the experimenter with one hand over the sternum. The bird was held for 10 s and then the hand was slowly removed. The duration of tonic immobility was recorded up to 600 sec. If the bird stood up within 30 sec. after the hand was removed from its sternum, new attempts to induce tonic immobility were made with up to three attempts per bird was allowed (for more information see Fogelholm et al., 2019). The birds were tested after sexual maturity at ∼170 days of age, with 7 days between each trial.

### Hypothalamus RNA isolation and microarrays

The Hypothalamus was selected for this analysis due to its pivotal role in the hypothalamic-pituitary-adrenal (HPA) axis and known effects on anxiety-related behaviour (File et al., 2000; McNaughton and Corr, 2004; Kallen et al., 2008), as well as its involvement in the regulation of social behaviour and sociality (for more information see Johnsson et al., 2016 and 2018). Immediately after being culled the hypothalamus was dissected out and snap frozen in liquid nitrogen, prior to storage at −80°. RNA was isolated with Ambion TRI Reagent (Life Technologies), according to the supplier’s protocol. Reverse transcription was performed with the Fermentas Revert Aid Premium First-Strand cDNA Kit and oligo-(dT)_18_ primers, followed by second-strand cDNA synthesis according to the supplier’s protocol (Thermo Fisher Scientific). All samples were quality checked on a Bioanalyzer chip (Agilent), and all had an RNA integrity number (RIN) value >8.1. Microarray hybridization and scanning was performed by NimbleGen Services (Reykjavik, Iceland).

#### Genotyping, QTL, and eQTL mapping

Using standard salt extraction on blood, DNA preparation was performed by Agowa (Berlin, Germany) and a total of 652 SNP were used to generate a map length of ∼9267.5 cM, with an average marker spacing of ∼16 cM (see Johnsson et al. 2015 for a full list of markers). QTL analysis was performed using the R/qtl software package (Broman et al. 2003) and interval mapping was performed using additive and additive + dominance models. Batch and sex were always included in the QTL behavioural analysis as fixed effects, whilst a principle component to account for population structure was included as a covariate. A sex-interaction effect was added, when significant, to account for a particular QTL varying between the sexes. Digenic epistatic analysis was performed according to Broman and Sen (2009) guidelines and a global model incorporated standard main effects, sex interactions and epistasis was built. The most significant loci were added to the model first, followed by the less significant loci. eQTL mapping was performed on the cross using R/qtl, as has already been documented previously (Johnsson et al. 2016). A local, potentially cis-acting, eQTL (defined as a QTL that was located close to the target gene affected) was called if a signal was detected in the closest flanking markers to the gene in question, to a minimum of 100 cM around the gene (i.e., 50 cM upstream and downstream of the gene). A distance of 50 cM was used to ensure that at least two markers up and downstream from the gene location were selected to enable interval mapping to be performed. The trans-eQTL scan encompassed the whole genome and used a genome-wide empirical significance threshold. In total, 535 local eQTL and 99 trans-eQTL were identified previously.

#### Significance thresholds

Significance thresholds for the behavioural QTL analysis were calculated by permutation tests (Churchill and Doerge 1994; Doerge and Churchill 1996). A genome-wide 20% threshold was considered suggestive, with this being more conservative than the standard suggestive threshold (Lander and Kruglyak 1995), while a 5% genome-wide level was significant. The ∼5% significant threshold was LOD ∼4.4, while the suggestive threshold was ∼3.6. Confidence intervals for each QTL were calculated with a 1.8 LOD drop method (i.e., where the LOD score on either side of the peak decreases by 1.8 LOD), with such a threshold giving an accurate 95% confidence interval for an intercross type population (Manichaikul et al. 2006). The nearest marker to this 1.8 LOD decrease was then used to give the confidence intervals in megabases. Epistatic interactions were also assessed using a permutation threshold generated using R/qtl, with a 20% suggestive and 5% significant genome-wide threshold once again used. In the case of epistatic loci, the approximate average LOD significance threshold for pairs of loci were as follows (using the guidelines given in Broman and Sen (2009) full model ∼11, full vs. one ∼9, interactive ∼7, additive ∼7, additive vs. one ∼4.

### Candidate Gene Analysis

To identify putatively causal genes underlying intra-individual behavioural variability, candidate QTL were overlapped with eQTL detected in the same cross (see Johnsson 2016). Once these QTL and eQTL were overlapped, each significant eQTL that overlapped a QTL was correlated with the behavioural trait of the QTL to test for a significance. For each eQTL overlapping a behaviour QTL, a linear model was fitted with the behaviour trait as a response variable and the expression trait as predictor, including sex and batch as factors. Weight at 42 days was included for traits where weight was used as a covariate in the QTL analysis. Any that were significantly correlated were then used for the final causality analysis using NEO (see below). One issue with using this approach with this particular data set is that the behavioural QTL were based on up to 572 individuals, whereas the eQTL/expression phenotypes were available only for 129 individuals. Therefore, the network edge orienting (NEO) method for causality testing was applied only where the behavioural QTL that a gene was potentially causative to was detectable in the smaller data set (*n* = 129). This technique has previously been successfully used with this intercross to detect genes that were potentially causal for both open field and social reinstatement behaviour (Johnsson et al., 2016 and 2018).

### Network edge orientation (NEO) analysis

Causality analysis was performed using NEO software (Aten *et al*. 2008) to test whether the expression of correlational candidates was consistent with the transcript having a causal effect on the behavioural trait. Single-marker analysis was performed with NEO fitting a causal model (marker → expression trait → behaviour) and three other types (reactive, confounded, and collider). The models tested by NEO were 1) CAUSAL: Genotype modifies gene expression which in turn modifies behaviour (genotype → expression trait → behaviour). 2)

REACTIVE: Genotype modifies behaviour which in turn modifies the expression trait (genotype → behaviour → expression trait). 3) CONFOUNDED: Genotype modifies both the expression trait and the behaviour separately (expression trait ← genotype → behaviour). 4) COLLIDER (behaviour is the collider): Genotype and the expression trait both independently modify behaviour (genotype → behaviour ← expression trait). 5) COLLIDER: (expression is the collider): Genotype and behaviour both independently modify the expression trait (genotype → expression trait ← behaviour). The leo.nb score quantifies the relative probability of the causal model (the preferred model) to the model with the next best fit (of the four remaining). The NEO software evaluates the fit of the model with a *χ*^2^-test, a higher *P*-value indicating a better fit of the model. The best-fitting model is chosen based on the ratio of the *χ*^2^ *P*-value to the *P*-value of the next best model on a logarithmic scale (base 10), called local edge orienting against the next best model (leo.nb) scores. A positive leo.nb score indicates that the causal model fits better than any competing model. Aten *et al*. (2008) use a single-marker leo.nb score of 1, corresponding to a 10-fold higher *P*-value of the causal model, and a multiple genetic marker leo.nb.oca of 0.3 as their threshold. More recently, other authors have relaxed this to use a threshold of 0.3 for the leo.nb score (see Faber et al., 2009, Plaisier et al., 2009). They also suggest that users inspect the *P*-value of the causal model to ensure the fit is good (in this case meaning the model *P*-value should be non-significant if the causal model fits the best). In effect this *P*-value is the probability of *another* model fitting the observed data. For each gene, we report the leo.nb score and *P*-value of the causal model. Leo.nb scores of 0.3 or more were considered suggestive, whilst leo.nb scores of 1.0 or more were considered significant.

### Global gene expression

Further analysis was performed for each intra-individual behavioural variance score whereby global gene expression (each gene, in turn, covering all 36,000 probesets) was correlated with behavioural variance. We used a linear model with the within-individual variance as response variable and expression, sex, and batch as predictors. To control for the large numbers of probes tested, we performed a permutation test. For any given behaviour, the behavioural variable was permuted, with this permuted phenotype then tested against all 36,000 probesets. The top 0.1% value was then retained from this permutation. A total of 500 permutations were performed for each behavioural measure.

### Data availability

Microarray data for the chicken hypothalamus tissue are available at E-MTAB-3154 in ArrayExpress. Full genotype and phenotype data are available on figshare with the following doi:XXXXXX

## RESULTS

### Within-individual behavioural variability is consistent within tests and contexts

Correlations of the different intra-individual behavioural variability scores show that individuals were consistently predictable or unpredictable within the SR-test and OF-test, (Table 1). For some behavioural traits, the magnitude of intra-individual variability was even more strongly positive correlated between behaviours in different test-situations, than behaviours within the same test situation (see table 1). These correlations were seen between IIV scores obtained from the OF-test and the SR-test (such as magnitude of IIV in ‘movement’ in the SR test and magnitude of IIV in ‘velocity’ or ‘time-spend-in-centre’ in the OF test, see supplementary material for list of measured behavioural traits). The same pattern could be seen in the correlations between average IIV scores obtained from the OF-test and SR-test and the individual IIV variability scores obtained from each trait (see table 1).The magnitude of IIV within the TI test was uncorrelated with virtually all the individual predictability measures.

**Table 1.**
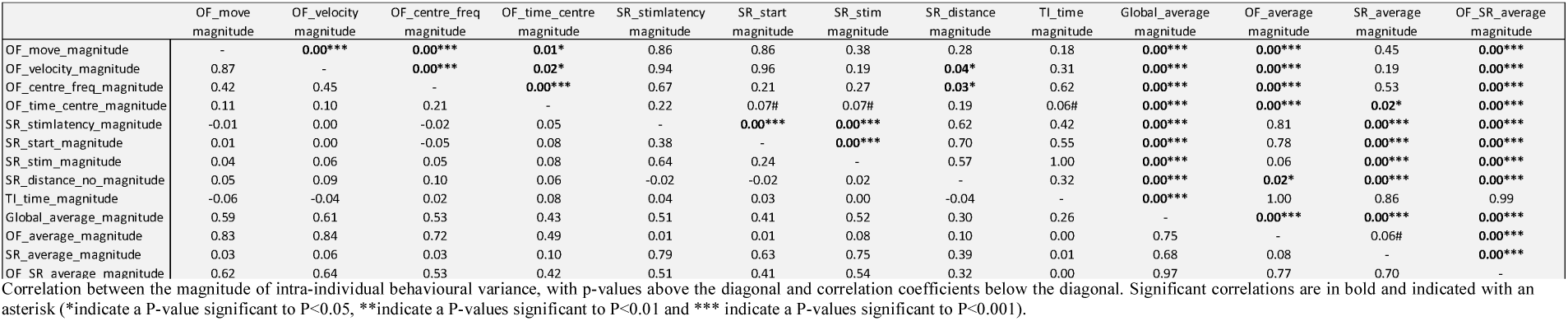
Intra-individual behavioural variance correlations.

|  | OF_move<br>magnitude | OF_velocity<br>magnitude | OF_centre_freq<br>magnitude | OF_time_centre<br>magnitude | SR_stimlatency<br>magnitude | SR_start<br>magnitude | SR_stim<br>magnitude | SR_distance<br>magnitude | TI_time<br>magnitude | Global_average<br>magnitude | OF_average<br>magnitude | SR_average<br>magnitude | OF_SR_a verage<br>magnitude |
| --- | --- | --- | --- | --- | --- | --- | --- | --- | --- | --- | --- | --- | --- |
| OF_move_magnitude | - | **0.00\*\*\*** | **0.00\*\*\*** | **0.01\*** | 0.86 | 0.86 | 0.38 | 0.28 | 0.18 | **0.00\*\*\*** | **0.00\*\*\*** | 0.45 | **0.00\*\*\*** |
| OF_velocity_magnitude | 0.87 | - | **0.00\*\*\*** | **0.02\*** | 0.94 | 0.96 | 0.19 | **0.04\*** | 0.31 | **0.00\*\*\*** | **0.00\*\*\*** | 0.19 | **0.00\*\*\*** |
| OF_centre_freq_magnitude | 0.42 | 0.45 | - | **0.00\*\*\*** | 0.67 | 0.21 | 0.27 | **0.03\*** | 0.62 | **0.00\*\*\*** | **0.00\*\*\*** | 0.53 | **0.00\*\*\*** |
| OF_time_centre_magnitude | 0.11 | 0.10 | 0.21 | - | 0.22 | 0.07# | 0.07# | 0.19 | 0.06# | **0.00\*\*\*** | **0.00\*\*\*** | **0.02\*** | **0.00\*\*\*** |
| SR_stimlatency_magnitude | -0.01 | 0.00 | -0.02 | 0.05 | - | **0.00\*\*\*** | **0.00\*\*\*** | 0.62 | 0.42 | **0.00\*\*\*** | 0.81 | **0.00\*\*\*** | **0.00\*\*\*** |
| SR_start_magnitude | 0.01 | 0.00 | -0.05 | 0.08 | 0.38 | - | **0.00\*\*\*** | 0.70 | 0.55 | **0.00\*\*\*** | 0.78 | **0.00\*\*\*** | **0.00\*\*\*** |
| SR_stim_magnitude | 0.04 | 0.06 | 0.05 | 0.08 | 0.64 | 0.24 | - | 0.57 | 1.00 | **0.00\*\*\*** | 0.06 | **0.00\*\*\*** | **0.00\*\*\*** |
| SR_distance_no_magnitude | 0.05 | 0.09 | 0.10 | 0.06 | -0.02 | -0.02 | 0.02 | - | 0.32 | **0.00\*\*\*** | **0.02\*** | **0.00\*\*\*** | **0.00\*\*\*** |
| TI_time_magnitude | -0.06 | -0.04 | 0.02 | 0.08 | 0.04 | 0.03 | 0.00 | -0.04 | - | **0.00\*\*\*** | 1.00 | 0.86 | 0.99 |
| Global_average_magnitude | 0.59 | 0.61 | 0.53 | 0.43 | 0.51 | 0.41 | 0.52 | 0.30 | 0.26 | - | **0.00\*\*\*** | **0.00\*\*\*** | **0.00\*\*\*** |
| OF_average_magnitude | 0.83 | 0.84 | 0.72 | 0.49 | 0.01 | 0.01 | 0.08 | 0.10 | 0.00 | 0.75 | - | 0.06# | **0.00\*\*\*** |
| SR_average_magnitude | 0.03 | 0.06 | 0.03 | 0.10 | 0.79 | 0.63 | 0.75 | 0.39 | 0.01 | 0.68 | 0.08 | - | **0.00\*\*\*** |
| OF SR average magnitude | 0.62 | 0.64 | 0.53 | 0.42 | 0.51 | 0.41 | 0.54 | 0.32 | 0.00 | 0.97 | 0.77 | 0.70 | - |

Correlation between the magnitude of intra-individual behavioural variance, with p-values above the diagonal and correlation coefficients below the diagonal. Significant correlations are in bold and indicated with an asterisk (\*indicate a P-value significant to P<0.05, \*\*indicate a P-values significant to P<0.01 and \*\*\* indicate a P-values significant to P<0.001).

### Unique genetic architecture of intra-individual behavioural variation

The chicken population used in this study was an advanced intercross based on Red Junglefowl (the wild progenitor of the modern domestic chicken) and domestic White leghorn chickens (for more information see Johnsson et al., 2016), and have previously been used to identify candidate genes for social behaviour (Johnsson et al., 2018), anxiety behaviour (Johnsson et al., 2016) and brain composition (Henriksen et al., 2016) in chickens. By integrating the IIV traits as part of a QTL mapping experiment using the advanced intercross framework, a total of 18 different QTL (quantitative trait locus associated with IIV scores) were identified. These included eleven QTL for the magnitude of IIV within the OF-test, two QTL for the magnitude of IIV within SR-test, three QTL for the magnitude of IIV for the TI-test and one QTL for ‘global’ magnitude of IIV (an overall measure of IIV using all the behavioural traits – see figure 2). Overall, the genetic architecture for IIV was largely unique as compared to the genetic architecture for the standard behavioural measures they were based on (see figure 2, and Johnsson et al., 2016, 2018, Fogelholm et al., 2019), even when correcting for the mean value of the behavioural traits (see supplementary table 2). Only three QTL, located on chromosome 10 (from 126-130cM) and one QTL on chromosome 24 (located at 77cM) overlapped any of the previously detected QTL for behavioural scores in the OF and SR (see figure 2, and Johnsson et al., 2016, and Johnsson et al., 2018). Furthermore, it should be noted that the above IIV QTLs did not overlap the behavioural QTL from the mean behavioural traits they were based on (see figure 2). These finding demonstrate for the first time that behavioural IIV is a trait which is under separate genetic control and adds proof to the concept of behavioural IIV being a trait in and of itself.

**Figure 2.**
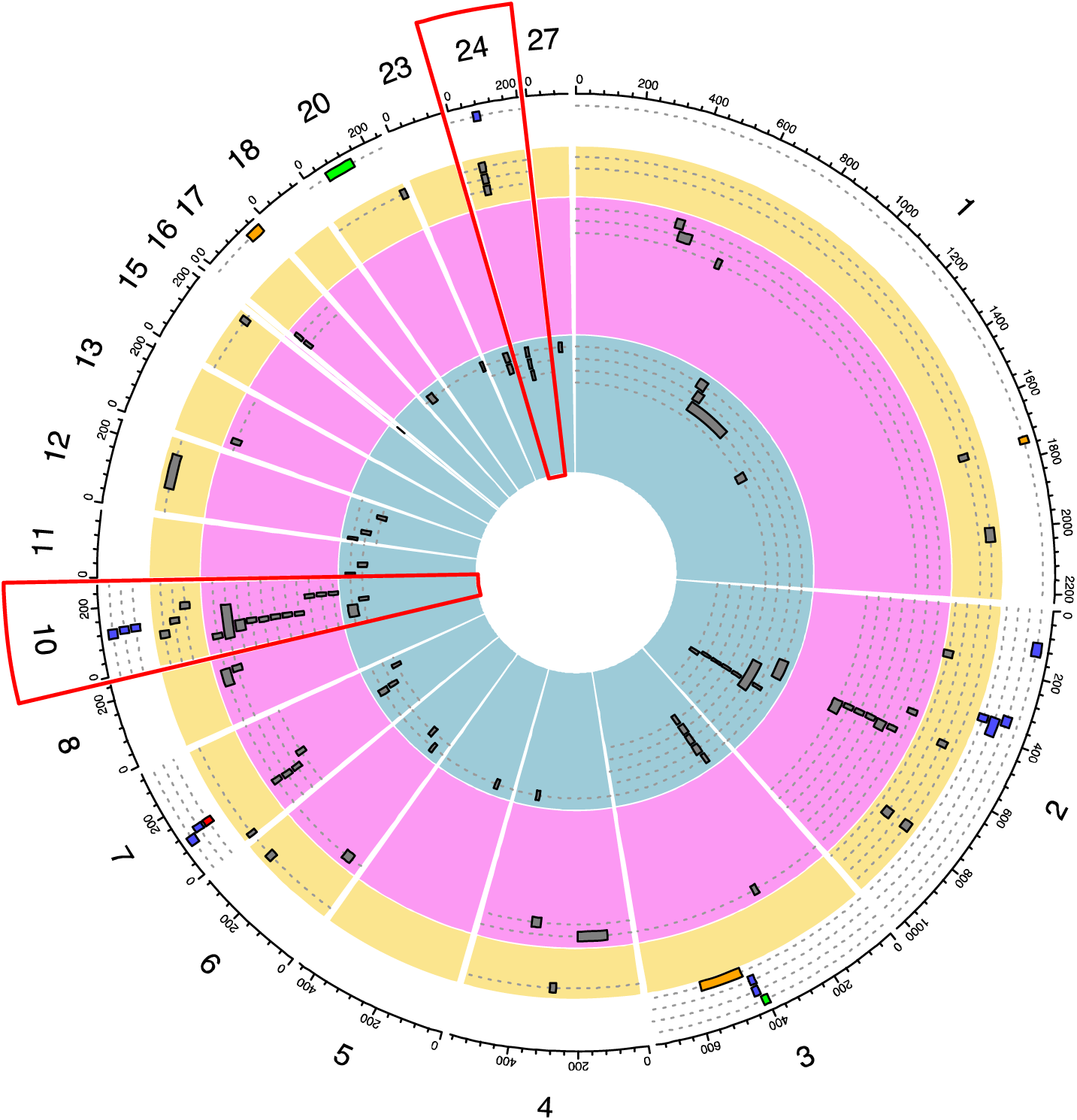
Unique genetic architecture of IIV in behavior. Circle plot showing the location of QTLs affecting magnitude of unpredictability (outer white circle) and mean behavioral trait values within the TI-test (yellow circle), OF-test (pink circle) and SR-test (inner blue circle). Predictability QTLs are based on the TI-test (orange bars), OF-test (blue bars), SR-test (green bars) and all tree test (global, red bars). Overlap between QTLs affecting the trait predictability and anxiety trait mean value are indicated with red boxed.

Interestingly, the QTL for the magnitude of behavioural IIV were generally in the direction of the domestic White Leghorn allele (8 strongly for White Leghorn allele, 4 strongly for Red Junglefowl allele) meaning that the majority of the alleles associated with increased behavioural IIV are derived from the domesticated allele.

### Candidate Genes for Intra-individual behavioural variance

To identify potential candidate genes that are involved in forming different IIV phenotypes, we combined quantitative trait locus (QTL) and expression QTL (eQTL) analyses of the brains from the advanced intercross. Intra-individual behavioural variance QTL were overlapped with eQTL, then the relevant behavioural IIV trait was correlated with the overlapped expression trait to identify those that not only overlapped but also exhibited a significant correlation (as used in Johnsson et al., 2016, and Johnsson et al., 2018). This initial analysis led to the identification of 10 putative genes underlying phenotypic differences in behavioural IIV, with these genes then taken to the next step of causality analysis (see table 2).

**Table 2.** Candidate genes and causality scores.

| Trait - behavioral IIV | QTL chromosome and position (cM) | gene | SINGLE MODEL |  | COMBINED MODEL |  | NEO |  |
| --- | --- | --- | --- | --- | --- | --- | --- | --- |
|  |  |  | gene expression (P-value) | genotype (P-value) | gene expression (P-value) | genotype (P-value) | LEO score | model P-value |
| OF_distance_moved_magnitude | chr 3 - 419 | MAP7 | 0.05 | 0.02 | 0.04 | 0.03 | 0.36 | 0.09 |
| OF_distance_moved_magnitude | chr 2 - 351 | SFRP4 | 0.05 | NS | NA | NA | 0.29 | 0.92 |
| OF_velocity_magnitude | chr 3 - 417 | MAP7 | 0.02 | 0.12 | 0.02 | 0.11 | 0.23 | 0.06 |
| OF_velocity_magnitude | chr 2 - 350 | SFRP4 | 0.02 | 0.05 | 0.02 | 0.05 | 0.5 | 0.84 |
| OF_velocity_magnitude | chr 10 - 130 | X603600179F1 (LOC100136711) | 0.04 | NS | NA | NA | 0.51 | 0.72 |
| OF_average_magnitude | chr 2 - 35 | X603862030F1(TJP1) | 0.07 | NS | NA | NA | -0.33 | 0.36 |
| OF_average_magnitude | chr 2 - 354 | SFRP4 | 0.03 | NS | NA | NA | 0.16 | 0.07 |
| SR_stimulus_zone_magnitude | chr 3 - 408 | X603599288F1(unknown) | 0.007 | 0.06/0.03 | 0.016 | 0.05/0.02 | 0.53 | 0.57 |
| SR_stimulus_zone_magnitude | chr 3 - 408 | X603600684F1(GALNT2) | 0.04 | 0.06/0.03 | 0.02 | 0.05/0.02 | 0.27 | 0.31 |
| TI_time_magnitude | chr 1 - 1750 | ITGBL1 | 0.005 | 0.03 | 0.005 | 0.14 | 0.51 | 0.33 |
| TI_time_magnitude | chr 1 - 1750 | ABHD13 | 0.05 | 0.03 | 0.04 | 0.09 | -0.11 | 0.17 |
| TI_time_magnitude | chr 3 - 564 | ENPP1 | 0.02 | NS | NA | NA | 0.34 | 0.26 |
| global average magnitude | chr 7 - 122 | X603601250F1(C2orf42) | 0.04 | 0.04 | 0.03 | 0.1 | 0.25* | <0.001* |

The intra-individual behavioral variance (IIV) QTL location (chromosome) and position (in cM) is provided. P-values are given for the significance of the gene expression and genotype (QTL) models, as predicting IIV. The single model p-values show these effects individually, whilst the combined model fits both gene expression and genotype simultaneously. The NEO causality score (LEO score) and NEO model P-values for assessing causality are also provided.

Since correlations alone are not enough to indicate the direction of the relationship between the candidate genes and behavioural, and support causality, we then utilized the network edge orienting (NEO) package (Aten et al., 2008), to infer causality between genotype, gene expression and behavioural IIV (Aten et al. 2008). The NEO package uses structural equation modelling to test the fit of five models each explaining a potential relationship between genotype (based on genetic marker), gene expression and behavioural IIV. The analysis produces a leo.nb score and a model p-value (see methods). The leo.nb score is defined as the log10 ratio of the causal model p-value to the next best model of the four remaining (all of which suggest a non-causal relationship between the behaviour and candidate gene). Therefore, a leo.nb score of 0.3 indicates that the causal model is the best fit, with twice as high model p-value than the next best model. In NEO and structural equation modelling in general, a small model p-value (e.g., p < 0.05) indicates a poor model fit. Our NEO analysis found 6 of the candidate genes (*Novel gene/X603599288F1, ITGBL1, SFRP4, X603600179F1/LOC100136711, MAP7, ENPP1*) passed the threshold of 0.3 for being suggestive, and are therefore potentially causative to IIV behaviours (see table 2). The NEO analysis also showed support for the gene C7H2oRF47 as controlling the magnitude of ‘global’ IIV (this model was done with multiple markers, which has a recommended threshold of 0.3, see Aten et al., 2008), with the model almost reaching the more stringent threshold suggested by Aten and colleagues.

Four of the candidate genes had previously been identified as having some bearing on “behaviour” (*MAP7, SRP4, ENPP1, LOC100136711*) or neuronal development (*MAP7*), while one had no previous evidence of functionality *(ITGBL1*) and one was a novel gene (*X603599288F1*).

## DISCUSSION

We find that intra-individual variation in behaviour is replicable between traits in the same behavioural test and to some extent between traits in different behavioural tests, although of similar contexts. These cross-test correlations demonstrate that individuals show consistency in their level of behavioural predictability both within and across test-type. While we might have expected that our measures of intra-individual variation in behaviour would be repeatable across traits measured within the same behavioural test, either because of correlated measurement error, or because they all respond in the same way to the same (unmeasured) environment gradient, positive correlations between intra-individual variation in behavioural traits across different types of assay are not auto-correlated, and therefore suggest that intra-individual variation in behaviour is a consistent trait (at least to some degree).

The genetic architecture for IIV was largely unique as compared to the genetic architecture for the standard averaged behavioural measures they were based on (Johnsson et al., 2016, 2018, Fogelholm et al., 2019). Of the total 18 QTL we identified to underlie between individual variation in IIV, only 3 QTL overlapped any of the previously detected QTL for the behavioural scores IIV were based on. However, the IIV QTL did not overlap the behavioural QTL from the mean behavioural traits they were based on. Our study thereby shows that within-individual variance in behaviour has a direct genetic basis which is largely unique compared to the genetic architecture for the standard behavioural measures they are based on.

We identified 6 candidate genes underlying intra-individual variation in behaviour. Some of these genes have previously been shown to be involved in natural behavioural variation and neural development, *SFRP4* and *LOC100136711* (the probeset was previously annotated as *LOC770352*, Johnssson et al., 2016) have, for example, been linked to anxiety in chicken (Johnsson et al., 2016), demonstrating some overlap in the genetic architecture of intra-individual variation in behaviour based on anxiety related traits and anxiety itself. Another of the candidate genes *ENPP1* has been linked to learning and memory abilities in mice, while *MAP7* has been shown to play a critical role in the developmental regulation of neural axon branch maturation (*MAP7*, Tymanskyj et al., 2017) and is also involved in schizophrenia (*MAP7*, Torri et al., 2010; Venkatasubramanian, 2015). Additional analysis whereby global gene expression was correlated with the magnitude of each IIV-trait (see supplementary table 1), highlighted the gene *CLDN5* as being significantly correlated at an experiment-wide level with the magnitude of three separate intra-individual variation in behaviour traits across the SR-test and OF-tests. Like *MAP7*, this gene has also previously been linked to Schizophrenia and highlights a potential link between schizophrenia and intra-individual variation in behaviour and the genes that underlie these traits. Increased within-individual behavioural variability characterizes the performance of people with schizophrenia and has been used for early diagnosis of this disorder (for review see McDonald et al., 2006).

The intra-individual behavioural variance we measure in our population, could be due to an innate difference in predictability between individuals, or alternatively could be due to inter-individual differences in adjustment to changes in novelty, such as habituation (decreased responsiveness to a stimulus with repeated presentation) or sensitization (increased responsiveness to a stimulus with repeated presentation, Dingemanse et al., 2012). Domestic ducks, for example, habituate faster than wild Mallard ducks (Desforges and Wood-Gush, 1975) and we find more domesticated alleles associated with intra-individual behavioural variation suggesting that the domesticated genotype produces an individual that is either more unpredictable in its behaviour or habituates faster. Domesticated chickens differ considerably in their behavioural responses from their wild progeny the Red Junglefowl, especially in anxiety related tests, such as the ones used in this study (Campler et al., 2009) and have also developed a brain that differs both in size and composition from their wild progeny (Henriksen et al., 2016). It is therefore not surprising that they might also differ in their behavioural predictability, perhaps due to altered selection pressure on this behavioural trait during domestication compared to the wild.

Intra-individual variation in the tonic immobility test did not correlate with the within-individual variation observed in the other test situations. One possible explanation could be that individuals were adult when measured in the TI test and chicks when measured in the OF and SR test. Age has previously been shown to affect intra-individual behavioural variation in Red Jungle fowls (Favati et al., 2016) perhaps because the cumulative experience of the environment leads to increasing consistency with age or because the neural circuitry has matured (MacDonald et al., 2006). Yet, the predictability of explorative behaviour in Great tits was shown to be unaffected by age (Dingemanse et al., 2002), and a large meta-analysis across animal species (by Belle et al., 2009) found no difference in the repeatability of behaviour between juveniles and adults. Another, explanation could be that the TI-test situation is different from the two other tests used in this study. In the TI-test a test-person physically restrains the bird to induce fear whereas in the SR-test and OF-test the test-person stays out of sight. Tonic immobility might therefore elicit a more direct fear response to a “predator” whereas the SR-test and OF-test measures an animal’s anxiety levels when feeling exposed and alone. This would indicate that intra-individual variation in behaviour is trait specific, at least to some extent. It would therefore be interesting in future studies to explore intra-individual variation in behaviour across more direct fear inducing tests, to see if the magnitude of intra-individual behavioural variation in the TI-test is test or trait-specific.

In conclusion, this study demonstrates that within-individual behavioural variation, like other behavioural traits, shows consistent differences between individuals and genotypes, even when animals are tested and reared in the same environment. This kind of behavioural variability between individuals has its own unique genetic architecture, separate from the behavioural traits it is based upon and therefore represents an important axis of consistent behavioural variation that evolution can act on. Our analysis also highlights six genes as putatively causative for intra-individual behavioural variation, four of which have been previously linked to behaviour and neural development.

## Acknowledgements

The research was carried out within the framework of the Swedish Centre of Excellence in Animal Welfare Science, and the Linköping University Neuro-network. SNP genotyping was performed by the Uppsala Sequencing Center. The project was supported by grants from the Carl Tryggers Stiftelse, Swedish Research Council (VR), the Linköping University Neuro-network and European Research Council (Consolidator grant FERALGEN 772874).

## Authors’ contributions

Conceived and designed the experiments: RH, MJ, DW. Performed the experiments: MJ, RH, AH, JF, DW. Analyzed the data: RH, DW, AH, RA. Wrote the paper: RH, MJ, DW, RA, AH, JF. All authors read and approved the final manuscript.

