## Supplementary material for "Intra-individual behavioural variability: a trait under genetic control": Supplmentary information

**ABBREVIATIONS**

SR	:	Social reinstatement

OF	:	Open field

TI	:	Tonic immobility

Magnitude	:	The absolute difference between the behavioural trait value obtained

during the first trial and second trial

SR_average_magnitude	:	The average magnitude for all traits measured within the SR-test

OF_average_magnitude	:	The average magnitude for all traits measured within the OF-test

Global_average_magnitude	:	The average magnitude for all traits measured within the SR-test, OF-

test and TI-test

**Social reinstatement test**

SR_stimlatency_magnitude	:	Latency to first enter the stimulus zone

SR_start_magnitude	:	Length of time in the start zone (the starting position of each bird,

farthest away from the stimulus zone)

SR_stim_magnitude	:	Length of time spent in the stimulus zone (adjacent to the three

conspecific birds)

SR_distance_no_magnitude	:	Total distance moved

**Open field test**

OF_move_magnitude	:	Total distance moved

OF_velocity_magnitude	:	Velocity

OF_centre_freq_magnitude	:	Proportion of time spent in the central zone

OF_time_centre_magnitude	:	Frequency (number of times) that the central zone was entered

**Tonic immobility test**

TI_time_magnitude	:	Duration of tonic immobility

**Supplementary Table 1**. A total of 59 genes were significantly or suggestively correlated at an experiment-wide level with the magnitude of a variety of the different behavioral predictability trait.

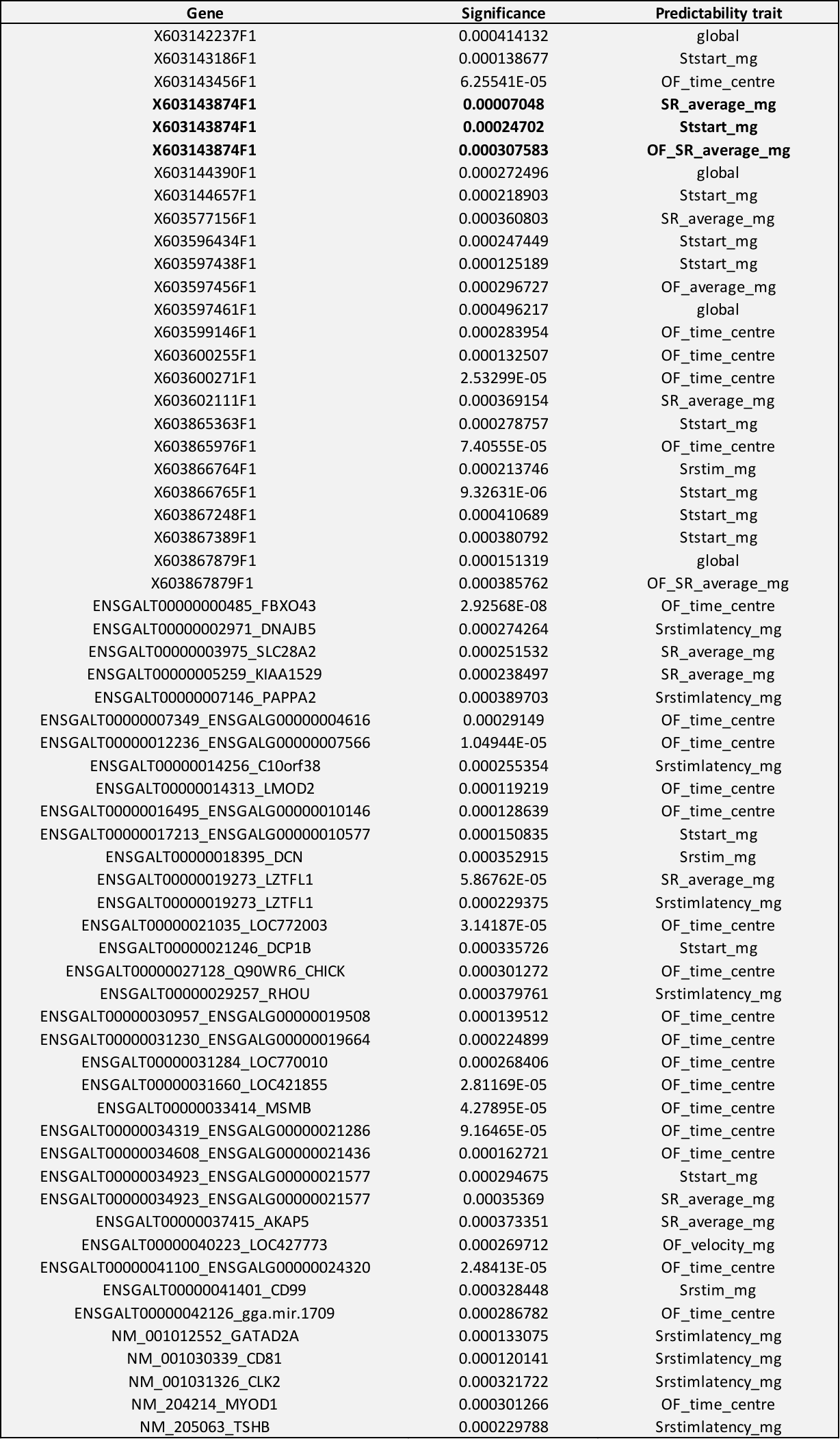

| **Supplementary Table 2**. QTL locations with average trait score included as a covariate | |
| --- | --- |
|  | **QTL chromosome** |
| **Trait - behavioral IIV** | **and position (cM)** |
| OF_move_mg | chr 3 - 419 |
| OF_move_mg | chr 2 - 353 |
| OF_move_mg | chr 10 - 130 |
| OF_move_mg | chr 7 - 121 |
| OF_time_mg | chr 2 - 121 |
| OF_time_mg | chr 24 - 77.1 |
| OF_velo_mg | chr 2 - 350 |
| OF_velo_mg | chr 10 - 132 |
| OF_velo_mg | ch 3 - 417 |
| OF_velo_mg | chr 7 - 109 |
| Srstim_mg | chr 3 - 408 |
| Srstim_mg | chr 20 - 112 |
| Titime_mg | chr 1 - 1734 |
| Titime_mg | chr 3 - 451 |
